# HAMRLNC: A Comprehensive Pipeline for High-throughput Analysis of Modified Ribonucleotides and Long Non-Coding Ribonucleic Acids

**DOI:** 10.1101/2025.05.28.656613

**Authors:** Chosen E. Obih, Jiatong Li, Giovanni Melandri, Duke Pauli, Eric Lyons, Andrew D. L. Nelson, Brian D. Gregory

## Abstract

As sequencing technologies advance and costs decline, there has been a surge in the application of RNA sequencing (RNA-seq) in understanding biological processes. In addition to the typical uses of RNA-seq for transcriptomics, gene annotation, novel gene discovery, and network analysis, these data can enable a deeper understanding of cellular processes through the identification of RNA modifications (epitranscriptome) and long non-coding RNAs (lncRNAs). To expedite discovery, we developed a portable, centralized computational pipeline for <u>H</u>igh-throughput <u>A</u>nnotation of <u>M</u>odified <u>R</u>ibonucleotides and <u>L</u>ong <u>N</u>on-<u>C</u>oding ribonucleic acids (HAMRLNC). HAMRLNC differs from existing methods by incorporating three workflows for quantifying transcript abundance, inferring RNA modifications, and lncRNA annotation using the same RNA-seq pre-processing and mapping steps. This facilitates reproducibility across multiple analyses and allows researchers to perform post-hoc analyses of archived sequencing data. In addition, we include novel analysis features to enable downstream visualization of annotated modified RNAs. HAMRLNC generates over a dozen well-defined and labeled figures as output, including gene ontology heatmaps, modification enrichment landscape, and modification clustering statistics.

**Availability and Implementation:** HAMRLNC is an open-source software, and the source code is available at https://github.com/bdgregory/HAMRLNC. The pipeline can be installed and used through a docker container (https://hub.docker.com/r/chosenobih/HAMRLNC/tags). HAMRLNC is also available as an app in the CyVerse Discovery Environment https://de.cyverse.org/.

**Supplementary information:** Supplementary data are available at BioRxiv online.

## 1 Introduction

RNA-seq is a valuable research method often used to gain insight into steady-state transcript abundance within a cell or tissue. Beyond its application in transcriptomics, RNA-Seq data are increasingly used for identifying lncRNAs and RNA modifications, both of which are emerging as important regulators of developmental processes and stress responses in plants and animals (Jha et al., 2020; Lu et al., 2020; Palos et al., 2023; Prall et al., 2023). Importantly, the same data that has been used to examine changes in transcript abundance in different contexts can also be used to identify novel transcripts (e.g., lncRNAs) or modification status on a set of transcripts. Previous approaches to utilize RNA-seq data for lncRNA and RNA modification identification, such as Evolinc and HAMR, have been quite successful in uncovering the complexity of eukaryotic transcriptomes (Nelson et al., 2017; Ryvkin et al., 2013). However, each of these pipelines was designed to run independently; they have different input requirements and data pre-processing steps and are even written in different coding languages. Given the importance of these hidden aspects of the transcriptome, what is needed is a unified framework that facilitates the analysis of all parts of the transcriptome at once, in both a scalable and user-friendly fashion. Here, we introduce a high-throughput computing pipeline, HAMRLNC, that automates the preprocessing steps of all RNA-seq data within an experiment to identify and quantify transcripts, modified RNA, and long non-coding ribonucleic acid molecules. HAMRLNC then plots patterns of modification deposition and transcript abundance across an experiment for the user to more easily interpret how the epitranscriptome is modulated within their data. HAMRLNC reduces the technical difficulty of these analyses, is more scalable, and increases reproducibility for comparative analyses of RNA-seq data.

## 2. Pipeline description

### 2.1 Input files

The pipeline minimally requires the following mandatory input files: a reference genome (FASTA), the companion reference genome annotation (GFF3), a metadata CSV table matching each FASTQ file to a sample group, and FASTQ file(s)/Sequence Read Archive (SRA) ID(s) or pre-mapped Binary Alignment (BAM) file(s).

### 2.2 Read trimming and quality control

For read trimming, fastp (v0.23.4, Chen et al., 2018), is used with default parameters to remove sequencing adapters and low-quality bases and FastQC (Andrew, 2010) is used for reads quality control assessment. Standard read trimmers have two modes of operation: single-end and paired-end, so this step applies to both single-end and paired-end sequencing data.

### 2.3 Read mapping and getting uniquely mapped reads

The trimmed reads are then mapped/aligned against the reference genome. STAR (v2.7.10a) (Dobin et al., 2013) is utilized for read mapping. If the -D flag is used to provide pre-mapped reads, the pipeline requires that the BAM files be mapped with an aligner and specifications that are suitable for the downstream analyses offered by HAMRLNC. Please see our documentation page for more details [https://github.com/bdgregory/HAMRLNC/wiki/Some-Notes-on-Pipeline-Design#mapping].

After the mapping step, SAMtools (v1.17) (Li et al., 2009) and Perl commands are used to extract uniquely mapped reads.

### 2.4 Preparing mapped reads and lncRNA annotation

First, Stringtie2 (v2.1.5) (Kovaka et al., 2019), is used to create a transcript assembly General Transfer Format (GTF) file from the uniquely mapped reads BAM file, and then the transcripts are merged to the reference annotation file. Next, we run GffCompare (v0.12.6) (Pertea et al., 2016) on the merged GTF and then filter out transcripts that are unstranded. The retained transcripts are further filtered using CPC2 (v1.0.1) (Kang et al., 2017) to remove transcripts with coding potential >= 0.5. Furthermore, all the intergenic transcripts are run through RFAM (v14.10) (Kalvari et al., 2018) to remove transcripts with similarity to known housekeeping RNAs (e.g., ribosomal RNAs). Lastly, we concatenate the final lncRNA GTF file to the reference annotation GTF file to produce an updated annotation file.

### 2.5 Preparing mapped reads and modified RNA annotation

GATK (v4.3.0.0) (McKenna et al., 2010) is used to process the uniquely mapped reads to resolve spliced alignments before running HAMR to annotate modified transcripts. First, read group IDs are added using picard AddOrReplaceReadGroups, the BAM files are reordered and indexed using SAMtools. Subsequently, GATK is used to resolve spliced alignments, then to resort the reads by coordinate. Finally, HAMR identifies RNA modification sites and annotates the RNA modification types using statistical and machine-learning approaches (Ryvkin et al., 2013).

### 2.6 Transcript abundance quantification

Transcript abundance quantification data is important for various downstream RNA-seq analyses. We implement featureCounts (Liao et al., 2014) for quantifying the relative abundance of transcripts from the uniquely mapped reads BAM file using the user-provided genome annotation file and the pipeline generated updated annotation file. To enable comparability, when the lncRNA annotation module is turned on, HAMRLNC runs a second transcript abundance step where the updated annotation file produced in the lncRNA annotation step is used for transcript abundance quantification.

**Fig. 1.**
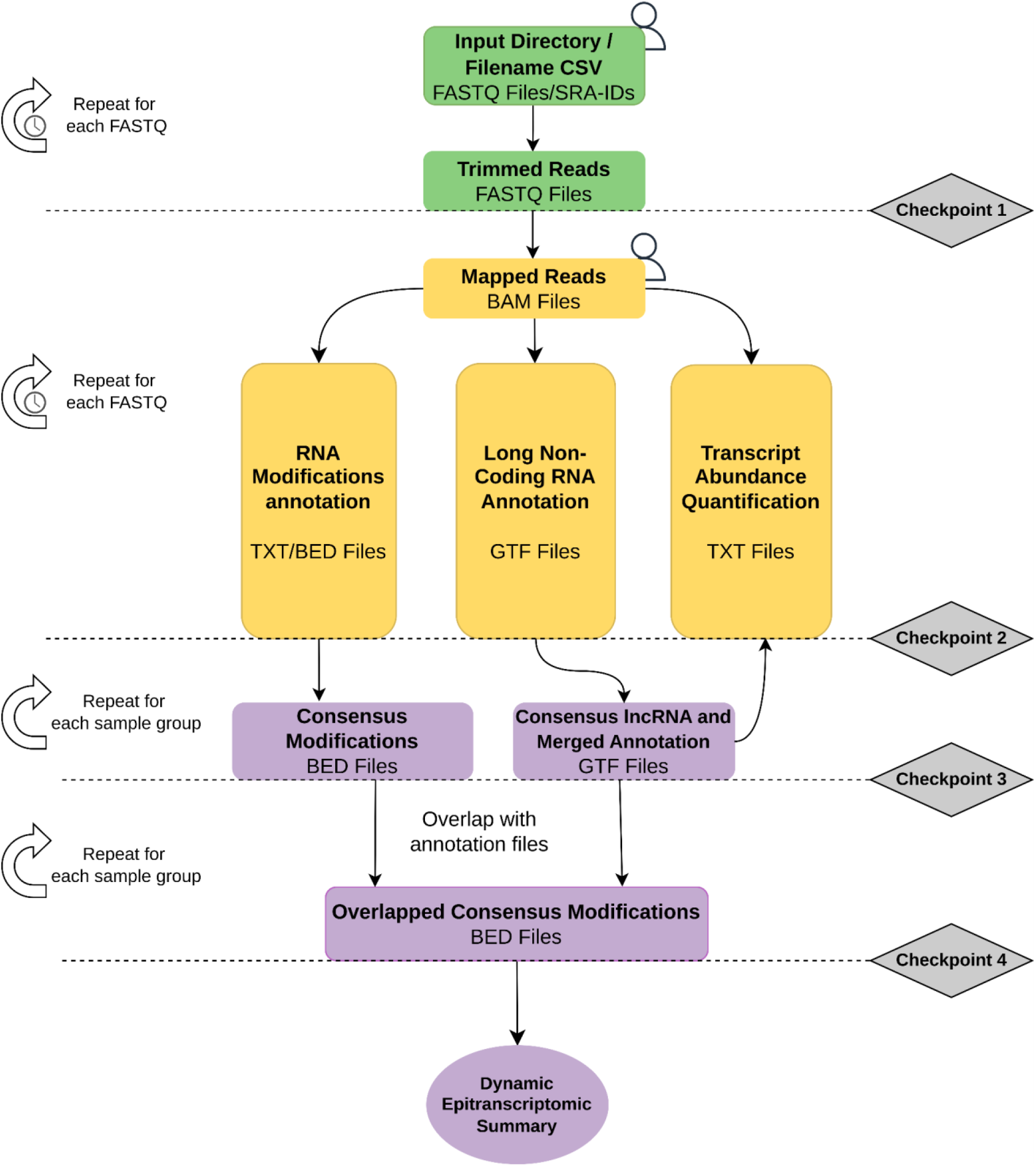
HAMRLNC workflow: This high-level summary shows the pipeline workflow, from raw RNA-seq data to dynamic epitranscriptomics summary. The input and quality control steps are in green, the pre-processing and core processing modules are in yellow, and the post-analyses steps are in purple. The checkpoints that enable restarting from the previous step in case of a failed run are in gray. Possible entry points of HAMRLNC are shown by the user icon. The clock icon indicates the time-consuming phases of the program.

### 2.7 RNA modification analyses

HAMRLNC produces several files/figures for visualization of predicted modified ribonucleotides. First, if the user provided multiple biological replicates per sample group, the union of pairwise intersections of the prediction set will be taken as the consensus. The consensus will then overlap with annotation libraries that HAMRLNC generates *de novo*, and the resulting predictions are concatenated in a CSV file, adapting a format similar to BED. HAMRLNC then automatically performs several analyses on this file, generating over a dozen well-defined and labeled figures as an output, including GO term heatmap (Supplementary Fig. 5), predicted enrichment landscape (Supplementary Fig. 4), modification clustering statistics (Supplementary Fig. 1, 2 & 3), etc. Users should refer to the pipeline’s documentation wiki page, https://github.com/bdgregory/HAMRLNC/wiki for each figure’s significance and underlying logic.

### 2.8 Pipeline output

All outputs of HAMRLNC are organized in corresponding subdirectories of the output directory. When run with all three core modules enabled, HAMRLNC produces twelve subdirectories in the output directory. Five subdirectories contain key intermediates like genome index files, trimmed FASTQ files, and bed files, which can be used in various downstream processing of the user’s choice. Three other subdirectories contain the raw output for each core functionality; one last subdirectory contains the visualizations and post analyses results. (Supplementary Fig. 1-5). We give the users the option to retain intermediate preprocessing files by activating the ‘-y’ flag.

## Supporting information

Pipeline visualizations figures and benchmarking

## Acknowledgements

We are grateful to all the members of the Epitranscriptomics research group made up of the Nelson lab at Boyce Thompson Institute, Gregory lab at the University of Pennsylvania and Lyons lab at the University of Arizona for their valuable feedback. We would like to thank Sarah Roberts and Michele Cosi (CyVerse) for their guidance in implementing HAMRLNC as an application on CyVerse Discovery Environment.

## References

Andrews, S. (2010). FastQC: A quality control tool for high throughput sequence data [Software]. Babraham Bioinformatics.

Chen, S., Zhou, Y., Chen, Y. & Gu, J. (2018). fastp: an ultra-fast all-in-one FASTQ preprocessor, Bioinformatics, 34(17), i884–i890.

Dobin, A., Davis, C. A., Schlesinger, F., Drenkow, J., Zaleski, C., Jha, S., Batut, P., Chaisson, M., & Gingeras, T. R. (2013). STAR: Ultrafast universal RNA-seq aligner. Bioinformatics, 29(1), 15–21.

Jha, U. C., Nayyar, H., Jha, R., Khurshid, M., Zhou, M., Mantri, N., & Siddique, K. H. M. (2020). Long non-coding RNAs: Emerging players regulating plant abiotic stress response and adaptation. BMC Plant Biology, 20(1), 466.

Kovaka, S., Zimin, A. V., Pertea, G. M., Razaghi, R., Salzberg, S. L., & Pertea, M. (2019). Transcriptome assembly from long-read RNA-seq alignments with StringTie2. Genome Biology, 20(1), 278.

Li, H., Handsaker, B., Wysoker, A., Fennell, T., Ruan, J., Homer, N., Marth, G., Abecasis, G., Durbin, R., & 1000 Genome Project Data Processing Subgroup. (2009). The Sequence Alignment/Map format and SAMtools. Bioinformatics, 25(16), 2078–2079.

Liao, Y., Smyth, G. K., & Shi, W. (2014). featureCounts: An efficient general purpose program for assigning sequence reads to genomic features. Bioinformatics, 30(7), 923–930.

Lu, L., Zhang, Y., He, Q., Qi, Z., Zhang, G., Xu, W., Yi, T., Wu, G., & Li, R. (2020). MTA, an RNA m6A Methyltransferase, Enhances Drought Tolerance by Regulating the Development of Trichomes and Roots in Poplar. International Journal of Molecular Sciences, 21(7), 2462.

McKenna, A., Hanna, M., Banks, E., Sivachenko, A., Cibulskis, K., Kernytsky, A., Garimella, K., Altshuler, D., Gabriel, S., Daly, M., & DePristo, M. A. (2010). The Genome Analysis Toolkit: A MapReduce framework for analyzing next-generation DNA sequencing data. Genome Research, 20(9), 1297–1303.

Nelson, A. D. L., Devisetty, U. K., Palos, K., Haug-Baltzell, A. K., Lyons, E., & Beilstein, M. A. (2017). Evolinc: A Tool for the Identification and Evolutionary Comparison of Long Intergenic Non-coding RNAs. Frontiers in Genetics, 8, 52.

Palos, K., Yu, L., Railey, C. E., Nelson Dittrich, A. C., & Nelson, A. D. L. (2023). Linking discoveries, mechanisms, and technologies to develop a clearer perspective on plant long noncoding RNAs. The Plant Cell, 35(6), 1762–1786.

Palos, K., Nelson Dittrich, A.C., Lyons, E.H., Gregory, B. D., & Nelson, A. D L. (2024) Comparative analyses suggest a link between mRNA splicing, stability, and RNA covalent modifications in flowering plants. BMC Plant Biology, 24, 768.

Pertea, M., Kim, D., Pertea, G. M., Leek, J. T., & Salzberg, S. L. (2016). Transcript-level expression analysis of RNA-seq experiments with HISAT, StringTie and Ballgown. Nature Protocols, 11(9), 1650–1667.

Prall, W., Ganguly, D. R., & Gregory, B. D. (2023). The covalent nucleotide modifications within plant mRNAs: What we know, how we find them, and what should be done in the future. The Plant Cell, 35(6), 1801–1816.

Ryvkin, P., Leung, Y. Y., Silverman, I. M., Childress, M., Valladares, O., Dragomir, I., Gregory, B. D., & Wang, L.-S. (2013). HAMR: High-throughput annotation of modified ribonucleotides. RNA, 19(12), 1684–1692.

Yu, X., Willmann, M. R., Vandivier, L. E., Trefely, S., Kramer, M. C., Shapiro, J., Guo, R., Lyons, E., Snyder, N. W., & Gregory, B. D. (2021). Messenger RNA 5′ NAD+ Capping Is a Dynamic Regulatory Epitranscriptome Mark That Is Required for Proper Response to Abscisic Acid in Arabidopsis. Developmental Cell, 56(1), 125-140.e6.

