## Supplementary material for "HAMRLNC: A Comprehensive Pipeline for High-throughput Analysis of Modified Ribonucleotides and Long Non-Coding Ribonucleic Acids": Pipeline visualizations figures and benchmarking

**Supplementary Data**

**Pipeline benchmarking**

The pipeline was benchmarked using a subset of single-end SRA data generated by Yu et al., 2021 with all three core processing flags (-k, -p, and -u) activated. Benchmarking was performed on an Intel_R_ Client Systems NUC10i5FNH equipped with an Intel® Core™ i5-10210U CPU @ 1.60GHz, with 8 cores, 64 GiB of memory, 2 TB of disk space, and Ubuntu 22.04.2 LTS as operating system. The internet speed was 970 Mbps. Additionally, to show the ability of the pipeline to reproduce results from published work, we re-processed a subset of paired-end data generated by Palos et al., 2024.

**Supplementary table**

**Supplementary table 1.** Information on benchmarking of the pipeline with a publicly available dataset

| Samples | Number of reads | Total run time (with genome index files provided) | Total run time (with genome index files not provided) |
|---|---|---|---|
| SRR10742163<br>SRR10742164<br>SRR10742165<br>SRR10742166 | 7,000,000<br>7,000,000<br>7,000,000<br>7,000,000 | 01:05 | 01:16 |

Run time, HH:MM

### Supplementary figures

### Supplementary Figure 1

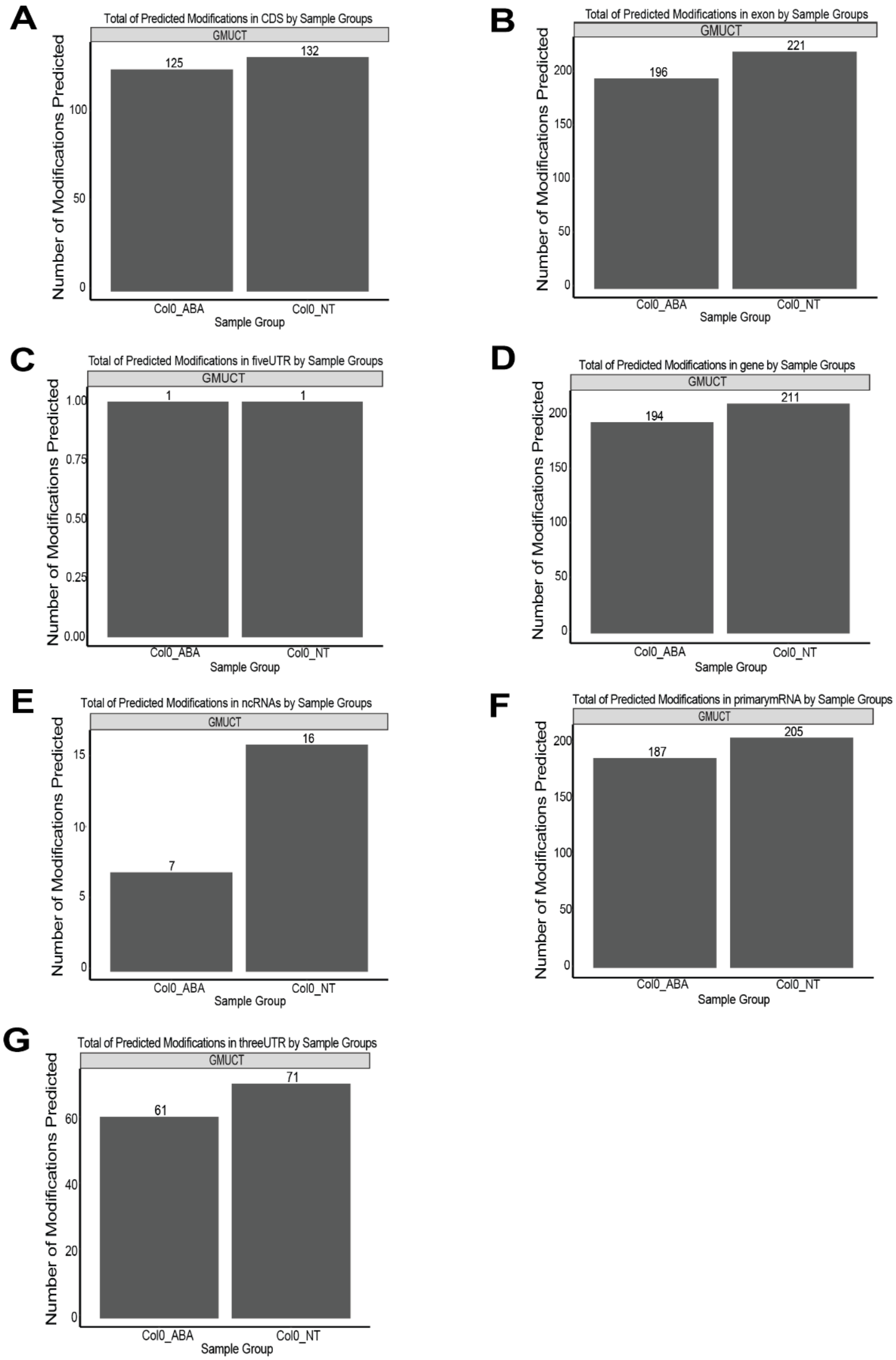


**Supplementary Fig. 1. (a-g)** Bar plots of the total abundance of HAMR predicted modifications by sample groups in CDS, exon, 5’ UTR, gene, ncRNA, primary mRNA, 3’ UTR regions

**Supplementary Figure 2**

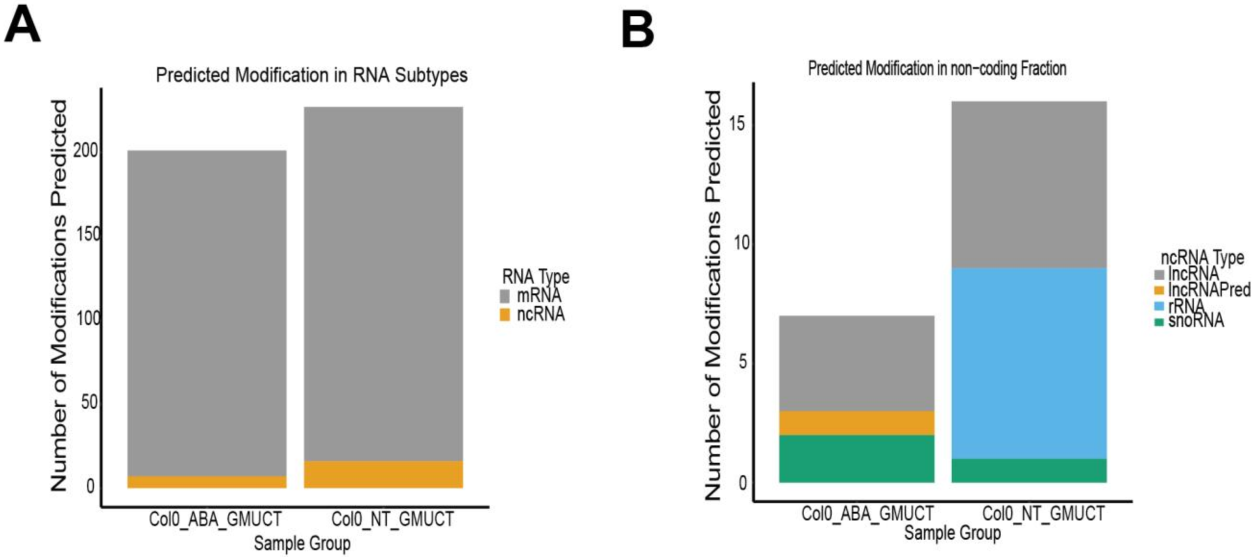


**Supplementary Fig. 2. (a-b)** Bar plots of HAMR predicted modification abundance located in different ncRNA types and RNA subtypes

Supplementary Figure 3

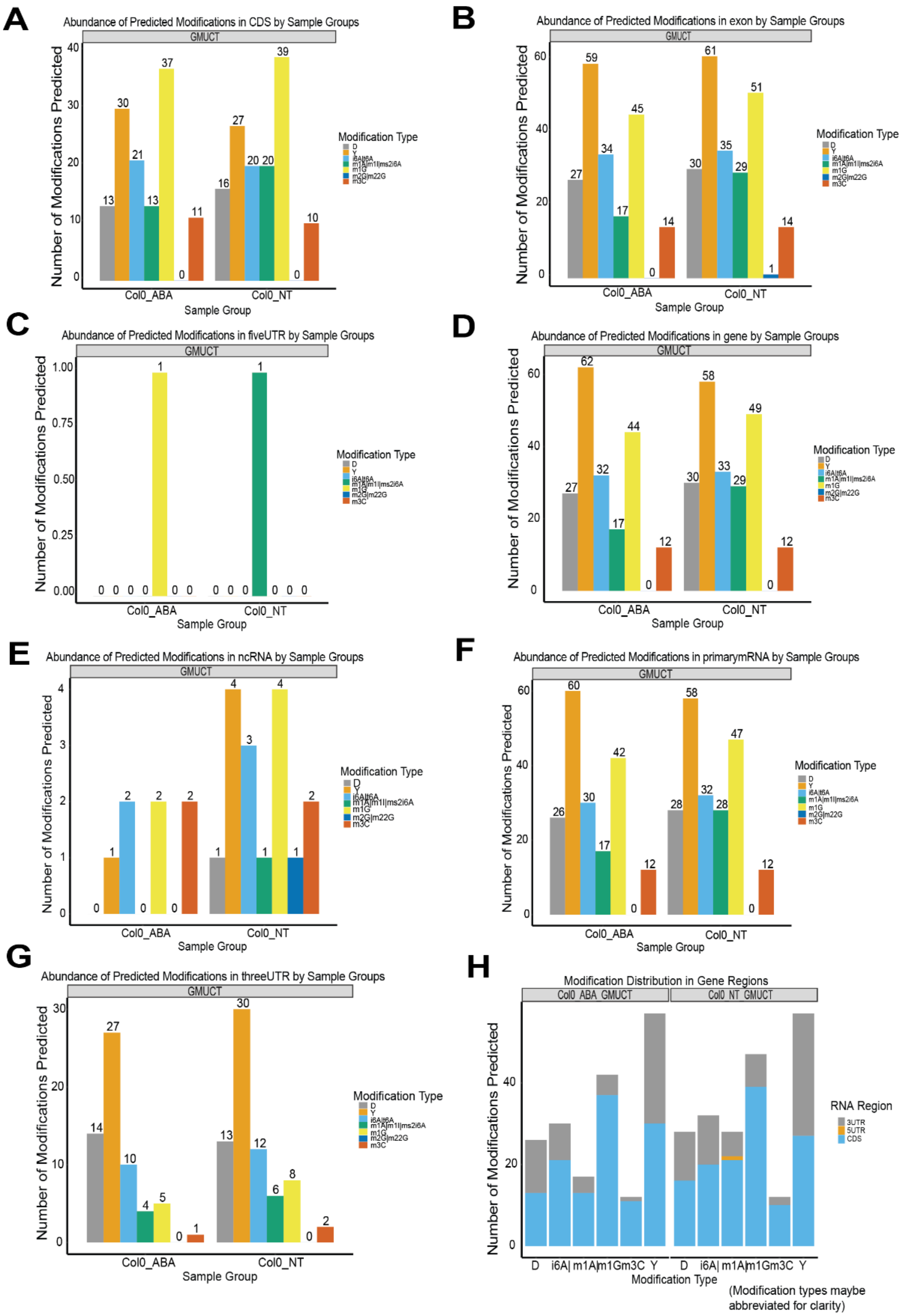


**Supplementary Fig. 3. (a-g)** Bar plots of the abundance of HAMR predicted modification classes by sample groups in CDS, exon, 5' UTR, gene, ncRNA, primary mRNA, 3' UTR regions. **(h)** Number of HAMR predicted modifications per gene region.

**Supplementary Figure 4**

A

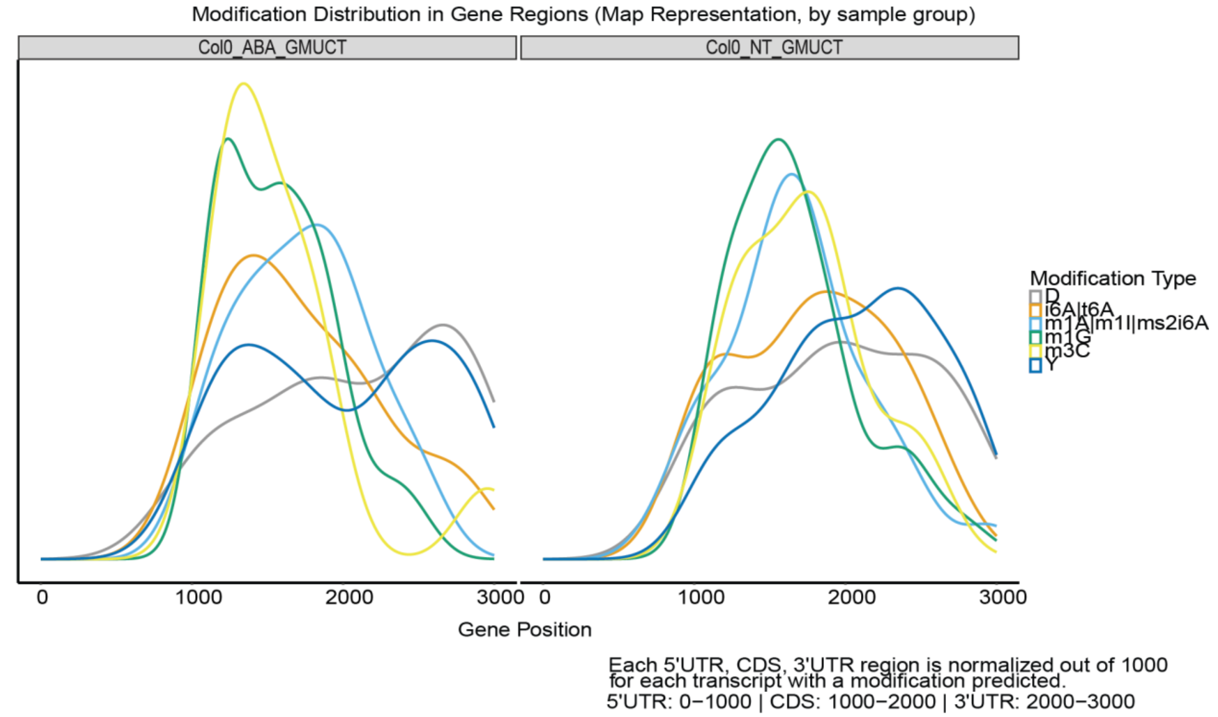


B

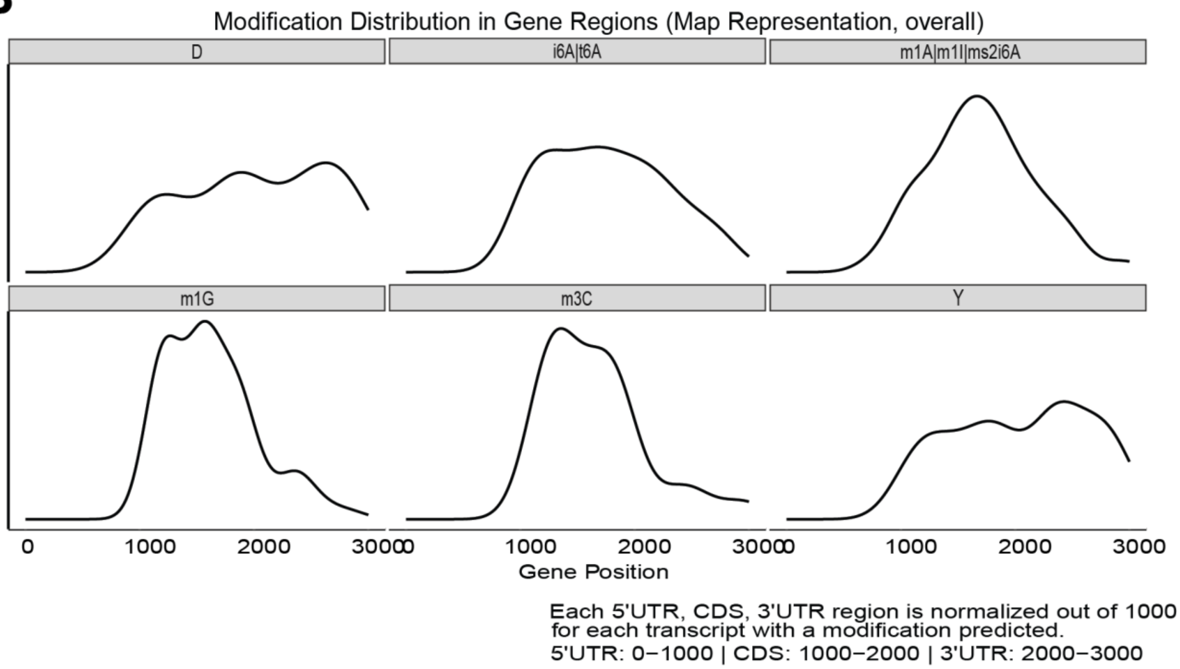


**Supplementary Fig. 4. (a)** Distribution of modification types in gene regions by sample groups. **(b)** Distribution of modification types in gene regions.

**Supplementary Figure 5**

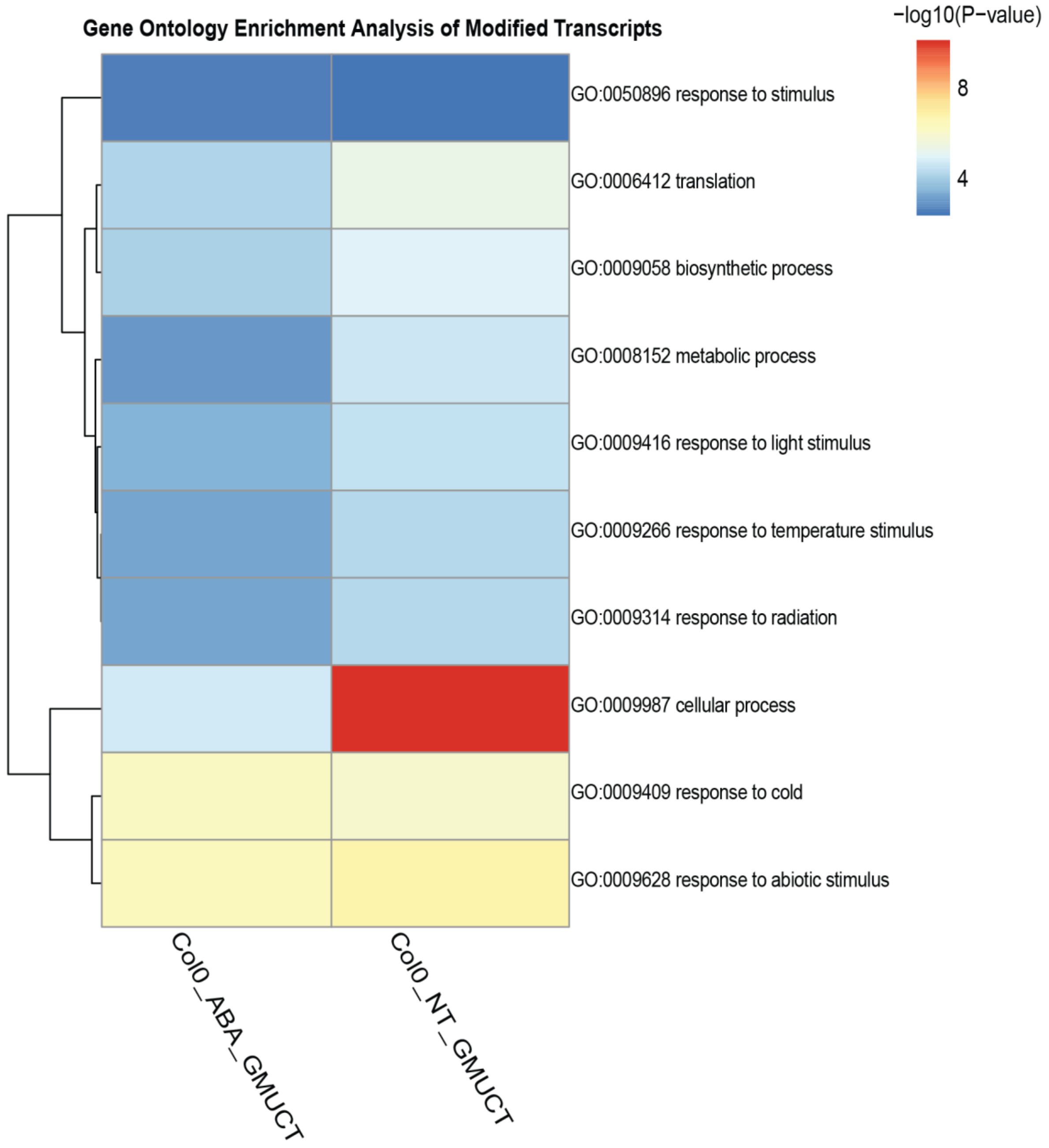


**Supplementary Fig. 5.** GO term heatmap and predicted enrichment landscape of modified transcripts.

**Supplementary Figure 6**

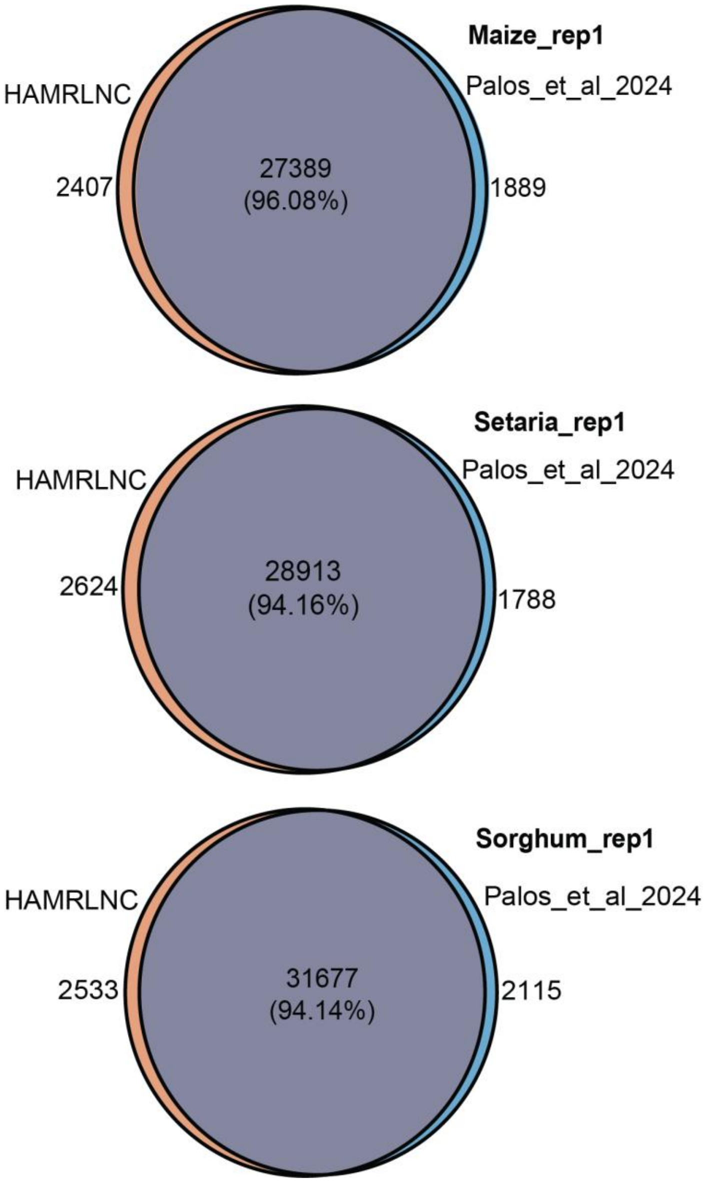


**Supplementary Fig. 6.** Venn diagram showing the overlap of modified sites (chromosomes and base pair positions) and predicted modification types reported by Palos et al., 2024 for three different grass species and HAMRLNC report.
